# Structural Variants in SARS-CoV-2 Occur at Template-Switching Hotspots

**DOI:** 10.1101/2020.09.01.278952

**Authors:** Brianna Chrisman, Kelley Paskov, Nate Stockham, Kevin Tabatabaei, Jae-Yoon Jung, Peter Washington, Maya Varma, Min Woo Sun, Sepideh Maleki, Dennis P. Wall

## Abstract

The evolutionary dynamics of SARS-CoV-2 have been carefully monitored since the COVID-19 pandemic began in December 2019, however, analysis has focused primarily on single nucleotide polymorphisms and largely ignored the role of structural variants (SVs) as well as recombination in SARS-CoV-2 evolution. Using sequences from the GISAID database, we catalogue over 100 insertions and deletions in the SARS-CoV-2 consensus sequences. We hypothesize that these indels are artifacts of imperfect homologous recombination between SARS-CoV-2 replicates, and provide four independent pieces of evidence. (1) The SVs from the GISAID consensus sequences are clustered at specific regions of the genome. (2) These regions are also enriched for 5’ and 3’ breakpoints in the transcription regulatory site (TRS) independent transcriptome, presumably sites of RNA-dependent RNA polymerase (RdRp) template-switching. (3) Within raw reads, these structural variant hotspots have cases of both high intra-host heterogeneity and intra-host homogeneity, suggesting that these structural variants are both consequences of *de novo* recombination events within a host and artifacts of previous recombination. (4) Within the RNA secondary structure, the indels occur in “arms” of the predicted folded RNA, suggesting that secondary structure may be a mechanism for TRS-independent template-switching in SARS-CoV-2 or other coronaviruses. These insights into the relationship between structural variation and recombination in SARS-CoV-2 can improve our reconstructions of the SARS-CoV-2 evolutionary history as well as our understanding of the process of RdRp template-switching in RNA viruses.

## INTRODUCTION

Researchers around the world are closely monitoring the evolutionary dynamics of SARS-CoV-2 (Severe acute respiratory syndrome coronavirus 2), the virus that causes COVID-19 (coronavirus disease 2019) and the source of the 2020 global pandemic. By studying the mutational patterns of viruses, we can better understand the selective pressures on different regions of the genome, robustness of a vaccine to future strains of a virus, and geographic dynamics of transmission.

Most evolutionary analysis begins with constructing a phylogenetic tree based on observed mutations or variants in various lineages of SARS-CoV-2. Most of these mutations are filtered to only single nucleotide polymorphisms (SNPs), as structural variants (SVs), particularly deletions, may be sequencing artifacts of low-quality reads or low-coverage genomic regions. This particular pipeline of analysis has two shortcomings. First, it ignores structural variants, despite their known role in viral evolution(27) and the importance of considering all types of mutations when building an accurate phylogenetic tree (20). Secondly, these phylogenetic trees are typically non-recurrent and do not take into account the possibility of recombination between viral lineages. Parallel research has been done to determine whether or not SARS-CoV-2 lineages have already recombined; however, the conclusions have been mixed (13, 26, 29). Not only does the relatively small number of mutations in the SARS-CoV-2 evolutionary history make it difficult to identify a clearly recombined lineage, additionally, the lack of publicly available raw reads makes it difficult to determine if seemingly recurrent mutations are due to recombination, site-specific hypermutability, or systematic sequencing error.

Recombination plays an integral role in the evolution of RNA viruses, including those implicated in recent epidemics: Comparative genomics studies suggest that SARS as well as a SARS-like coronavirus in bats have recombinant origins (9, 15), co-circulating and recombinant lineages of MERS-CoV were found in dromedary camels (22) and several studies hypothesize SARS-CoV-2 has a recombinant origin from bat coronaviruses, pangolin coronaviruses, or both(1, 14, 30, 31).

It is generally accepted that recombination in RNA viruses is via a copy-choice mechanism by which an RdRp switches template strands during negative strand synthesis, the first step of both sub-genomic transcription and full-genome replication in +ssRNA viruses (Fig. 1)(6, 19, 23). Outside of known transcription regulatory sites, what causes RdRp to disassociate and reassociate to a different template strand mid-transcription or replication is not well understood (25). An early study suggested that in the absence of natural selection, RNA virus recombination occurs entirely at random with respect to genome position (2), and is independent of RNA secondary structure or sequence. Successive studies have found secondary RNA structure motifs that lead to RdRp disassociation and subsequent recombination in RNA viruses (4, 8, 12, 21).

**Figure 1.**
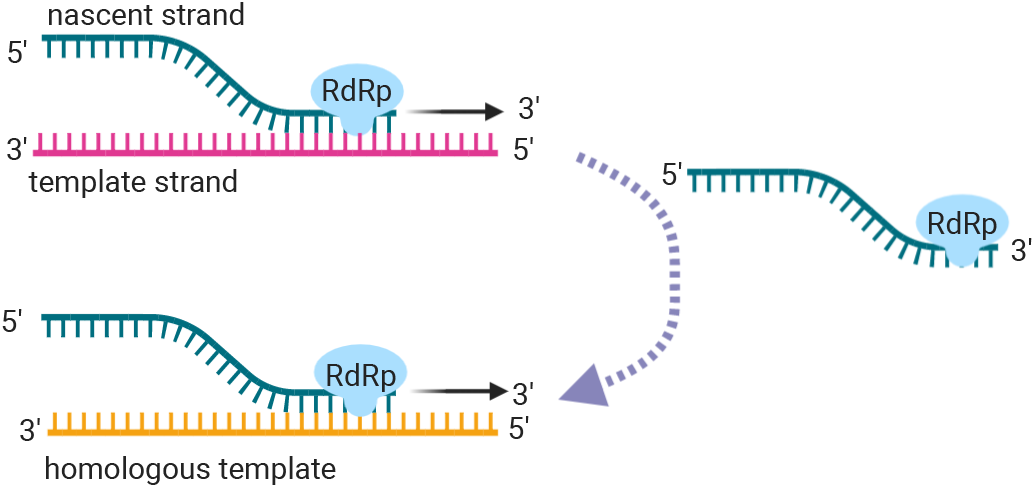
Copy-choice recombination is the presumed primary recombination mechanism for RNA viruses. During negative strand synthesis, the replication complex and the nascent strand disassociate from the template strand. From there, the replication complex can template-switch, or reassociate with a homologous or replicate template strand.

Using 16,419 GISAID sequences (24), we characterize over 100 deletions and insertions in the evolutionary history of SARS-Cov-2 as of early June 2020, and hypothesize that these indels are the result of imperfect homologous recombination. We offer four independent pieces of evidence that suggest this (Fig. 2). (1) We show that the indels in the consensus GISAID sequences are found in clusters across the genome. (2) Using long-read transcriptomic data (11), we show that these clusters correspond to regions of the genome that have high rates of TRS-independent polymerase jumping, hypothetically RdRp template-switching hotspots. (3) We show that these many of the structural variant hotspots show high rates of heterogeneity in the raw reads, suggesting that even sequences where the consensus sequence does not contain the structural variant may be undergoing *de novo* recombination at these sites. (4) Finally, we show that many of these indel clusters are found on “arms” within the predicted RNA secondary structure of SARS-CoV-2, suggesting that global RNA secondary structure may play a role in RdRp template-switching in SARS-CoV-2 and other coronaviruses.

**Figure 2.**
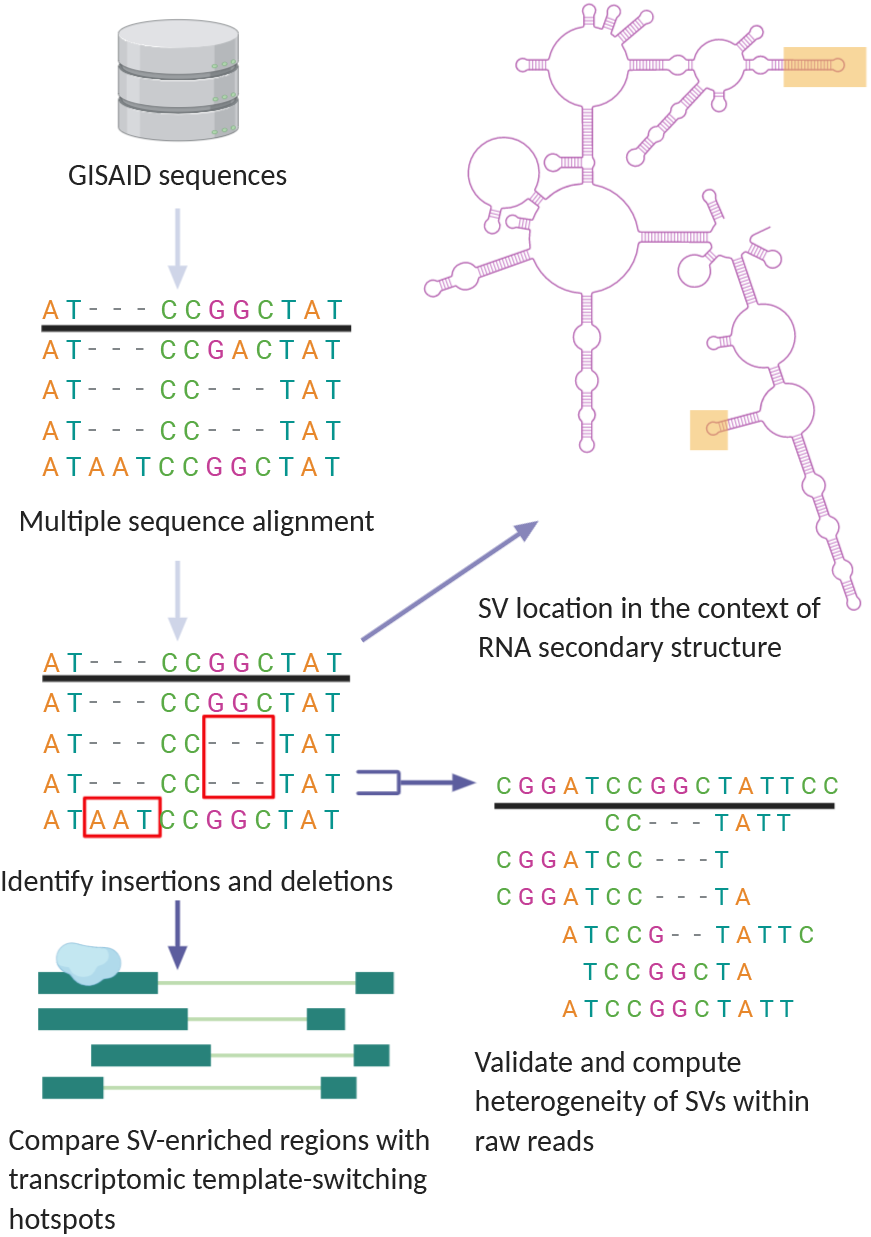
General pipeline of project: using GISAID sequences, we identified structural variants present in SARS-CoV-2 lineages. We compared the location of these SVs to regions of discontinuous transcription breakpoints, computed the heterogeneity of SVs using raw reads, and analyzed the SV locations with respect to the secondary RNA structure using a simulation of the folded SARS-CoV-2 RNA molecule.

## MATERIALS AND METHODS

### Data Access and Preparation

To obtain the SARS-CoV-2 consensus sequences, we accessed the GISAID sequences on June 3, 2020. We filtered to high-coverage full length (>29kb) sequences, where less than 20 bases were missing, totalling 16,419 sequences.

To obtain the raw reads, we accessed the NCBI SRA run browser on June 3, 2020. We found the accession numbers that corresponded SARS-CoV-2 reads that were full length, short reads from Illumina sequencing machines, and consisted of less than 1 billion total base calls (to speed up computation). We downloaded these using the NCBI’s `fastq-dump` API.

To compare the regions with enriched numbers of structural variants to the hypothesized SARS-CoV-2 template-switching hotspots, we used the deep sequencing long-read SARS-CoV-2 transcriptome data published by Kim *et al*(11). We used the reads from the Vero-infected cells, filtered to reads that aligned to the SARS-CoV-2 genome (`VeroInf24h.viral_genome.bam`).

### Identifying Structural Variants

We used `MAFFT10`(10) to perform multiple sequence alignment of the GISAID sequences with NC 045512.2 as the reference sequence. We locally realigned indel calls that were synonymous using an in-house python script. The subset of indels that were present in 2 or more sequences is shown in (Table), and the entire set of structural variants found among these sequences is shown in Table S1.

### Comparing with Template-Switching Hotspots in the Transcriptome

Kim *et al* (11) identified non-canonical subgenomic RNAs (sgRNAs) in the SARS-CoV-2 transcriptome characterized by large deletions in the middle of the transcript, presumably a result of the template-switching mechanism for discontinuous transcription. From the viral reads collected in the Kim *et al* study, we filtered to reads with a deletion of 100 bases or more relative to the reference genome and computed the locations of the 3’ and 5’ breakpoints. We compared the breakpoint hotspots to the location of the indels we identified in the GISAID sequences (Fig. 3).

**Figure 3.**
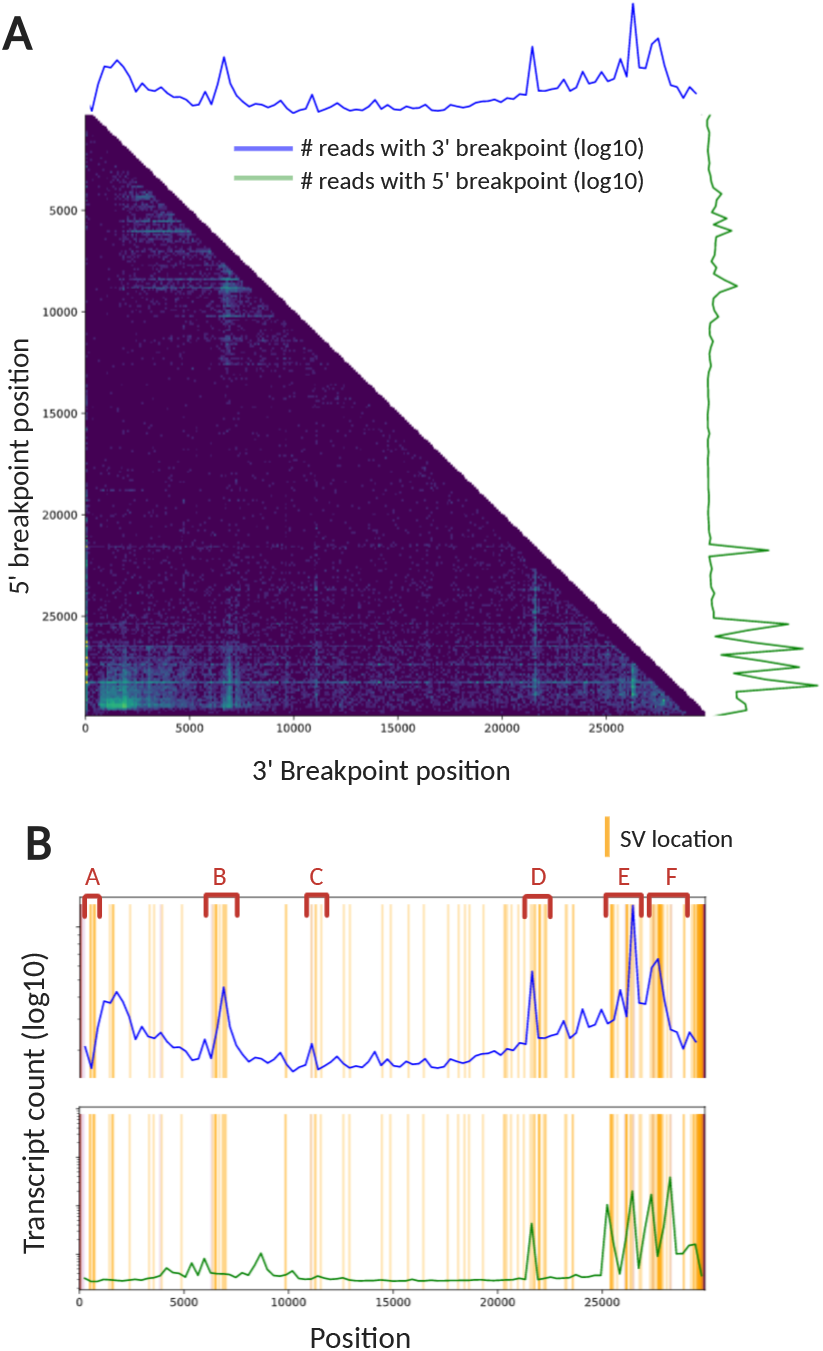
**(A)** Heatmap of 5’ breakpoint vs 3’ breakpoints from long-read TRS-independent discontinuous transcripts. Reads were considered TRS-independent discontinuous transcripts if the leader sequence was not near the transcription regulatory sequence and the aligned read contained a deletion of 100 or more bases. The heatmap and histograms are in bins of 100 bases. Colors in the heatmap and amplitude of blue and green lines are log-normalized. **(B)** The log-normalized number of discontinuous transcripts with given 5’ and 3’ breakpoints overlaid with the locations of the SVs (by 5’ start position)

### Structural Variant Heterogeneity from Raw Reads

We analyzed the indels in the context of the raw reads for two major reasons. First, we wished to validate that these structural variants were in fact true insertions or deletions, and not the result of sequencing error or low-coverage genomic regions. Secondly, we wished to measure intra-host heterogeneity at these sites, to determine whether these structural deletions might be the result of imperfect *de novo* recombination events or located in hypermutable regions, or whether they were inherited from the a viral lineage in the previous host.

We accessed NCBI’s SRA run browser on June 3, 2020 to download the `fastq` files for the full genomic sequences of SARS-CoV-2. We restricted to Illumina reads, as short reads have smaller error rates than long reads and are less prone to systematic sequencing types of errors (16). We quality filtered the reads using fastp (5), with a qualified quality phred cutoff of 20, an unqualified percent limit of 20, and a required length of 50. Using NC 045512.2 as the reference, we used bwa-mem (17) to align reads to the reference genome, following the standard paired for single-end read pipelines as appropriate. We marked and removed PCR duplicates using GATK’s MarkDuplicates. We used lofreq to quality score the indels, perform local realignment, and compute SV heterogeneity (28). For each sample, we considered a structural variant call to be homogeneous if the read depth was >50 and the alternate allele frequency was >95%. We considered a structural variant to be heterogeneous if the read depth was ¿50 and the alternate allele frequency was between 5% and 80%. We used an in-house python script to visualize the raw read alignment to the reference genome for a given sample and structural variant loci as shown in Fig. 5.

### Predicted RNA Secondary Structure

It is presumed that RdRp template switching is responsible for the discontinuous transcription and recombination in coronaviruses. While transcription regulatory sites (TRS) govern some of the leader-to-body fusion sites, little is known about what mechanisms are behind TRS-independent transcription and replication. We used RNAfold (18) with the default parameters, to generate an estimate of the secondary structure of the reference SARS-CoV-2 RNA genome. We visualized regions with high indel enrichment within the secondary structure using VARNA (7).

### Data and Code Availability

The full source code for the analysis can be found at https://github.com/briannachrisman/SARS-CoV-2SVs. The datasets used are all publicly available (or from GISAID), and access methods are described above.

## RESULTS

### SARS-CoV-2 lineages contain over 100 indels

Ignoring the error-prone and low-coverage 5’ and 3’ ends of the genome, we found 114 total indels between loci 100 and 29,800 (Table. S1).

Table shows all of the structural variants that were found in two or more sequences. Of these 40 SVs, all but 12 are deletions or insertions of multiples of 3-basepairs, and would not result in a frameshift. Many of the indels that would result in a frameshift occur downstream from loci 29500, after the stop codon of the last canonical open reading frame.

### Structural variants cluster at SARS-CoV-2 template-switching hotspots

The coronavirus transcriptome is characterized by discontinuous transcription events. During discontinuous transcription, RNA-dependent RNA polymerase (RdRp) ‘jumps’ from a 5’ breakpoint to a 3’ breakpoint. This discontinuous transcription may occur across a single genome of a virus or it may involve 2 copies of the RNA genome, with the RdRp switching from one template (leader) to another (body) mid-transcription. According to the prevailing model, leader 5’ breakpoints and body 3’ breakpoints occur at short motifs called transcription-regulatory sequences (TRSs) adjacent to open reading frames. In a deep sequencing study of the SARS-CoV-2 transcriptome, Kim *et. al*. found that there were many discontinuous transcription events not characterized by TRS (known as TRS-L-independent fusion), with both the 5’ and 3’ breakpoints clustered at specific regions of the genome. The mechanism behind TRS-L-independent fusion is not currently well understood.

Several genomic regions that we found to be enriched for deletions and insertions correspond to the regions the Kim *et al*. found to be enriched for 5’ and 3’ breakpoints. In Fig. 3, we note several regions of interest, where the genome was enriched for indels identified from the GISAID sequences and where the genome was enriched for either 5’ or 3’ breakpoints.

### Structural variants have intra-host heterogeneity

Using the raw reads, we found high rates of intra-host heterogeneity for the indels. Fig. 4 shows the rates of heterogeneity for SVs at each loci as computed from the raw reads. Many of the same regions of the genome enriched for 5’ and 3’ breakpoints in the transcriptome, particularly regions B, C, D, and F, also have high rates of heterogeneity for small deletions and insertions within the raw reads. Many indels have samples with high heterogeneity, but also samples with high homogeneity for a given structural variant call, as shown in Fig. 5. This suggests that structural variants may occur by either a recombination or mutation event in a previous host, resulting in high intra-host homogeneity, or from *de novo* recombination within a current host resulting in high intra-host heterogeneity.

**Figure 4.**
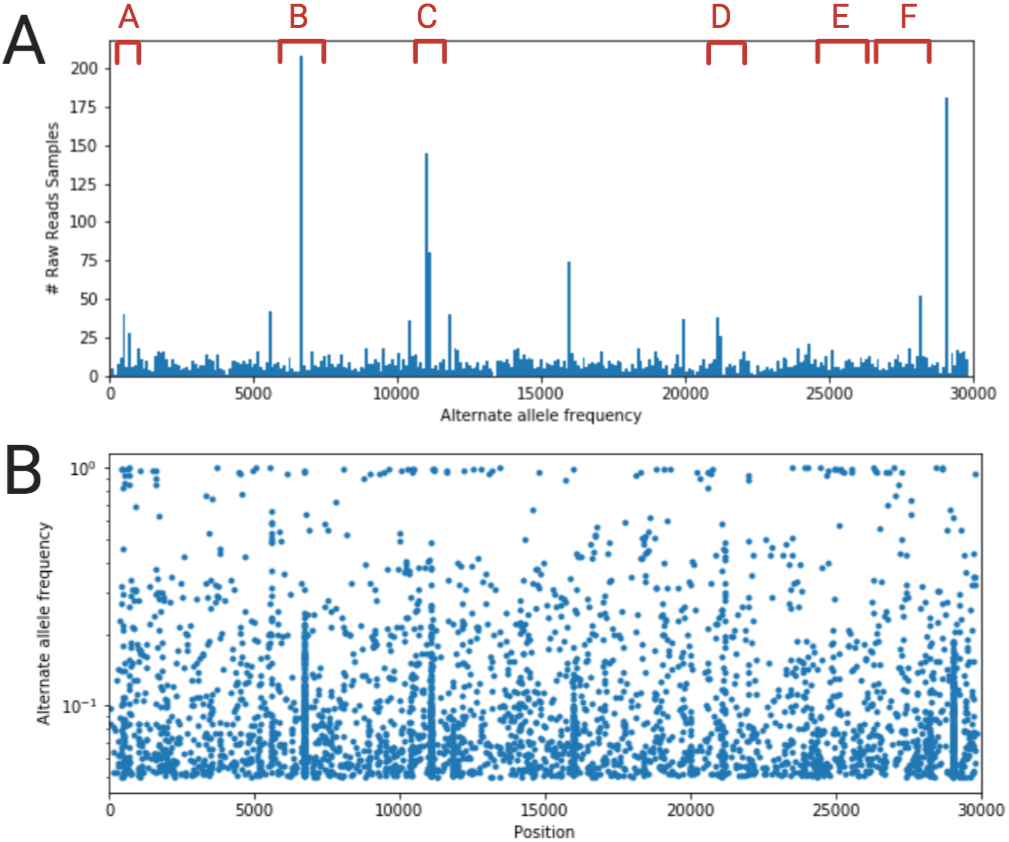
Rates of heterogeneity in raw reads. Filtered to raw reads with SV heterogeneity rates of >5% and read depth >50. **(A)** Histogram of number of samples with heterogeneity rate of >5% for an SV at given loci. **(B)** Alternate allele frequency vs SV location as computed from raw reads.

**Figure 5.**
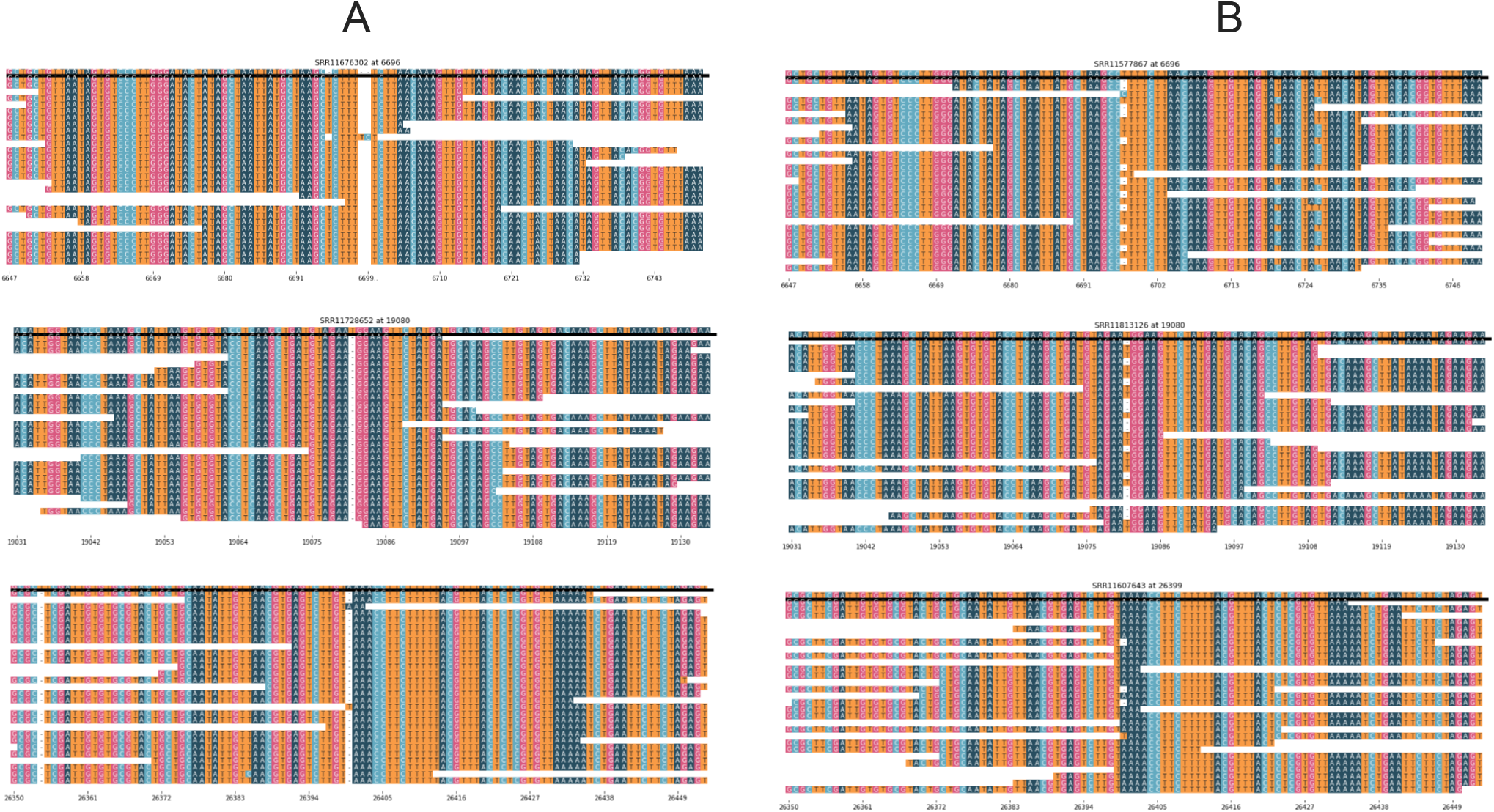
Raw read alignment for several common deletions Examples of the raw reads for SVs with strong homogeneity (¿95%)in one host and high heterogeneity (20-80%) in another host. **(A)** Examples of raw reads with a strong homogeneity at the SV, suggesting that the SV arose from an event in a previous host. Note that the small amount of heterogeneity can be attributed toward locally misaligned reads, where the SV was near the end of a read and thus difficult for lofreq to realign properly. **(B)** Examples of raw reads with intra-host heterogeneity at the SV, suggesting that these indels are *de novo* events.

### Structural variants cluster at arms in the secondary RNA structure

To see if there were any obvious structural motifs associated with SVs or hypothesized recombination hotspots, we simulated the secondary RNA structure of SARS-CoV-2 and analyzed the locations of the SV clusters (Fig 6 We see that structural variant clusters tend to be on “arms” of the folded RNA; that is highly accessible regions that are extended away from the RNA backbone. In particular, regions B, and D-F are located on the farthest extensions of the folded RNA molecule. Within the arms, many specific SVs appear to occur inside stem loop structures, although this is not always the case.

**Figure 6.**
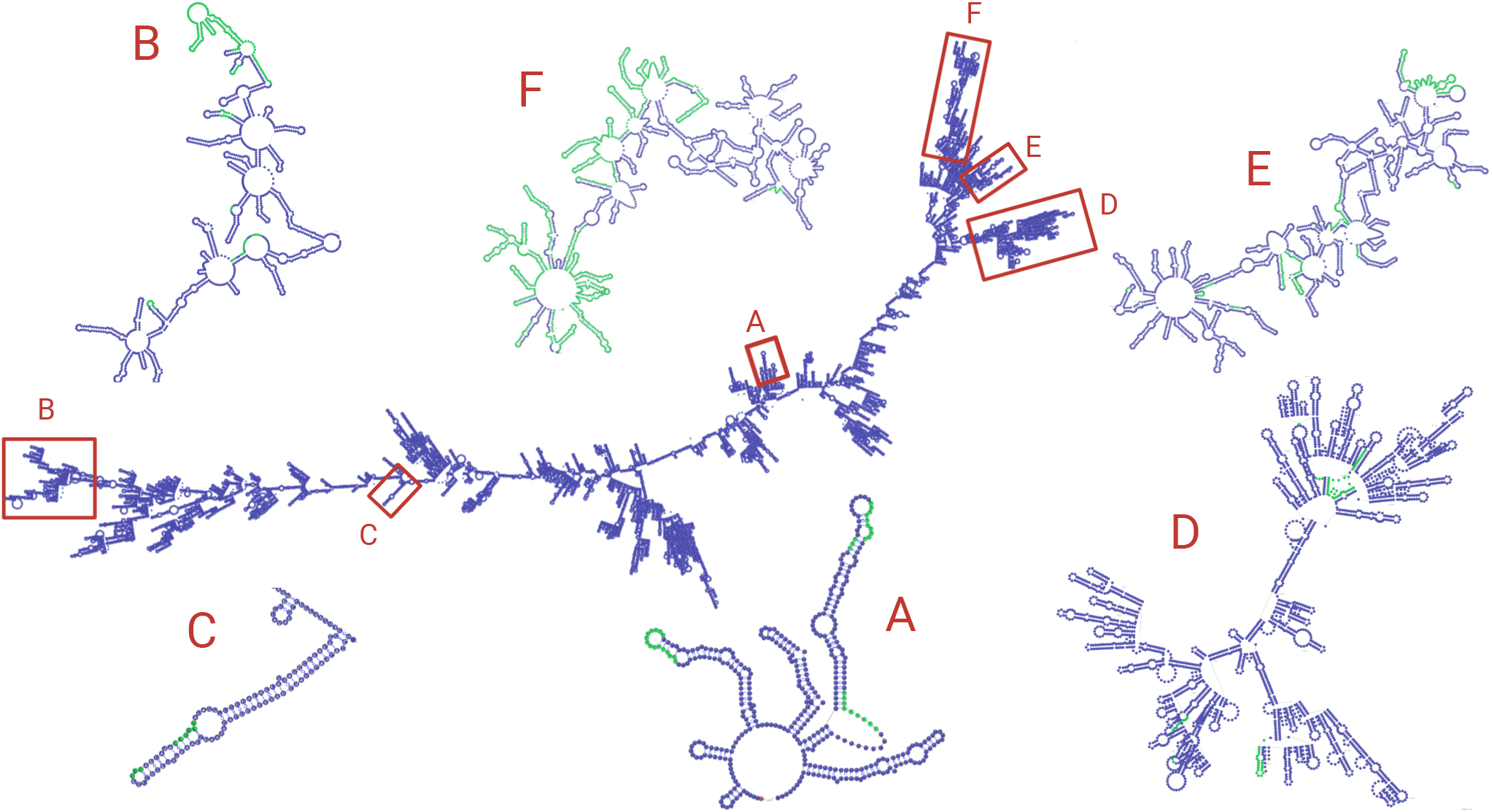
RNA structure was simulated using RNAfold and visualized with VARNA. The zoomed-in subsections of RNA are from the selected regions of both high SV and discontinuous transcription breakpoint enrichment. Green areas represent regions with an indel. Note that the structures zoomed-in subgenomic RNAs have been manually refined to avoid overlap of loops for easier visualization.

## DISCUSSION

We have catalogued over 100 structural variations in the SARS-CoV-2 genome, a type of mutation that has been largely ignored thus far in the analysis on SARS-CoV-2 evolutionary history. Via the GISAID consensus sequences, publicly available raw reads, long-read deep transcriptomic data, and simulated RNA structure, we show several independent pieces of evidence that suggest that these indels are artifacts of recombination, and that SARS-CoV-2 contains several recombination hotspots.

Interestingly, using sequence-based recombination detection approaches, previous studies have identified several of our hypothesized recombination hotspots as recombination breakpoints in SARS-CoV-2 and other related coronaviruses.

Lau et al. found evidence of the N and ORF8 proteins of SARS being acquired from recombination between horseshoe bat viruses. They identified recombination breakpoints at 20900, 26100, 27128, and 28635(15) - which correspond well to our SV hotspots D, E, and F. Hom *et al.* also identified a possible recombination breakpoint around 21495 in tracing SARS from a bat coronavirus(9), corresponding to SV-enriched region D. Lam *et al.* identified a possible recombination schema for SARS-CoV-2 from Malayan panglolin viruses and bat CoVs with breakpoints around 11000, 21000, 23000, 24000(14), which corresponds to C, D, and E SV-enriched regions. Analysis on the sarbecovirus recombinant origins of SARS-CoV-2 identified possible recombination breakpoints at 1684, 3046, 9237, 11885, 21753, 22773 and 24628.(3). 1684 is close to 1605, at which 345 sequences have a 3-bp deletion. The latter 4 breakpoints fall close to or within our identified SV-enriched region C, D, and E. We also see several deletions between 2500-3500 (possibly linked to the breakpoint hotspot at 3046), though we see no SVs within 500b of 9237.

Globally, regions enriched for SVs and transcriptional breakpoints appear to fall on “arms” of the simulated folded RNA molecule. We hypothesize the because these regions of the RNA molecule are extensions from the backbone, they are easily accessible and therefore the RdRp can “jump” between homologously aligned replicate molecules. We note that this is a crude representation of the secondary RNA structure; it ignores the interactions between genome and nucleocapsid, uses only the reference sequence and does not capture how mutations might change the folded RNA structure in different lineages, ignores psueodknots, and only shows the primary consensus fold. Taking into account these nuances, it seems likely the global structure of the full genome would still be a good estimate, but the local structures (as shown in the zoomed-in regions in Fig. 6) may not be reliable. Additional work needs to be done to determine if local sequence or structural motifs exist that guide RdRp disassociation.

There are several alternative explanations for these highly enriched regions of structural variation, but we believe that they are unsupported by the combined evidence in the GISAID sequences, raw reads, and transcriptome data. First of all, addressing the obvious possibility of systematic sequencing or alignment error, we see no signs in the raw read data that the indels are due to such error types. The indels occur in many Illumina samples, which are not prone to systematic sequencing errors, and many samples have nearly 100% homogeneous calls for a given indel.

Another theory is that perhaps structural variants occur at hypermutable sites within the genome, and it is either by chance that they appear to be clustered in several regions, or selective pressure weeds out structural variations in other areas of the genome. However, recall that these regions are also enriched for 5’ and 3’ breakpoints in the transcriptome, which we calculated by only considering reads with a deletion > 100 basepairs. If these sites are in fact hypermutable, then they are also hypermutable for larger structural variations as well; selective pressure would not be acting on the transcriptome in such a manner. It seems possible, however, that there might be additional template-switching hotspots that can be seen in the discontinuous transcriptome, but not in the regions of SV enrichment because selective pressure makes SARS-CoV-2 unable to handle SVs in this region. For example, there seems to be enrichment of 5’ end breakpoints in the discontinuous transcriptome between loci 8000 and 9000, however we found no SVs in that region (see Fig. 3); perhaps a SV in this region would result in a dysfunctional phenotype.

Finally, these indels might be the result of RdRp disassociating and reassociating from one location to another on the same strand of RNA, rather than from a template strand to a nascent strand of a viral replicate. This would mean that these SVs are not created from template switching between two separate viral strands, but from RdRp disassociating and reassociating on the same viral strand. This is possible; however it is likely that if an area is a hotspot for RdRp jumping within the same strand, it is consequently a hotspot for RdRp template switching between two different template strands. Recombination between two or more SARS-CoV-2 template strands could be verified experimentally by measuring recombination rates between mutant viral lineages, or computationally by finding a patient that has been co-infected by two different SARS-CoV-2 lineages with discernible mutations on either side of a recombination breakpoint. This computational verification may be difficult as it would require co-infection in a patient, the presence of both lineages within the same cell, recombination, and the recombinant lineage to make it into the sequencing reads.

We emphasize how valuable the raw or aligned reads are for better understanding of SARS-CoV-2 evolutionary dynamics. Although the consensus sequences such as those on GISAID provide some information about mutational patterns and evolutionary dynamics, there are several shortcomings in consensus sequences that raw reads can address. As we have shown, using raw reads we can quantify site-specific mutability. An estimate of per-site variation, both for SNPs and for SVs, is essential for building accurate phylogenetic trees, which can then be used to trace the spread of SARS-CoV-2 and identify recurrent mutations or sites under high selective pressure. Furthermore, as SARS-CoV-2 continues to spread and inevitably recombines with either itself in the form of a different lineage, another coronavirus, or another RNA molecule, the raw reads with provide a clearer understanding of recombination patterns than consensus sequencing can.

We therefore urge the scientific community to make their raw reads publicly available if possible. While there are possible privacy concerns with human DNA or RNA contamination in the data, most pipelines that generate a consensus sequence involve filtering our reads aligning to the human genome, thereby maintaining privacy and lowering barriers for open access to the scientific community.

In conclusion, we have catalogued over 100 indels present in the SARS-CoV-2 evolutionary history thus far and shown several independent pieces of evidence that these clusters of indels indicate recombination hotspots. An improved understanding of structural variation as well as recombination in coronaviruses will improve phylogenetic reconstructions of the evolutionary history of SARS-CoV-2 and other coronaviruses, and is one step closer to understanding the outstanding questions surrounding the RdRp template-switching mechanism in RNA viruses.

**Table 1.** Table of structural variants (Deletion - D, insertion - I) found in at least two sequences.

| Start Pos | Length | Type of SV | # Sequences | Countries |
| --- | --- | --- | --- | --- |
| 508 | 15 | D | 2 | USA |
| 510 | 9 | D | 6 | France, Scotland, England, USA, Australia |
| 515 | 6 | D | 11 | Bangladesh, England, USA, Belgium, Greece, Denmark |
| 669 | 3 | D | 9 | USA, India |
| 686 | 9 | D | 48 | France, Turkey, Iceland, England, USA, Portugal, Australia, Spain, Sweden, Israel, Belgium, Denmark, Netherlands, Canada, Saudi |
| 729 | 9 | D | 5 | Wuhan, Sichuan |
| 1431 | 3 | D | 2 | Yunnan, USA |
| 1605 | 3 | D | 341 | Scotland, Australia, Taiwan, Chile, New, Germany, Greece, Netherlands, Russia, USA, Latvia, Pakistan, NewZealand, Northern, France, Wales, Iceland, Lithuania, England, Portugal, Spain, Sweden, Finland, Belgium, Denmark, Canada |
| 3333 | 3 | D | 23 | Kazakhstan |
| 6501 | 3 | D | 2 | England |
| 6510 | 6 | D | 2 | India, Australia |
| 6518 | 6 | D | 2 | USA |
| 11074 | 3 | I | 17 | Switzerland, Jordan, Scotland, England, Portugal, United, Australia, Jamaica, Taiwan |
| 12620 | 3 | D | 2 | Netherlands |
| 14865 | 2 | D | 12 | Wuhan |
| 18412 | 1 | D | 6 | Wuhan |
| 20423 | 3 | D | 2 | Portugal, USA |
| 20965 | 1 | D | 4 | Wuhan |
| 21990 | 3 | D | 2 | Belgium, USA |
| 21991 | 3 | D | 12 | Slovenia, Jordan, India, England, USA, Netherlands, Saudi |
| 25532 | 3 | D | 2 | France, USA |
| 26159 | 2 | D | 3 | USA |
| 26351 | 6 | D | 3 | India |
| 27471 | 201 | D | 3 | Bangladesh |
| 27701 | 3 | D | 2 | England |
| 27792 | 3 | D | 2 | China, Kazakhstan |
| 27848 | 382 | D | 13 | Singapore |
| 27910 | 345 | D | 2 | Bangladesh |
| 28090 | 6 | D | 4 | Canada, USA, Iceland, Australia |
| 28254 | 1 | D | 6 | Wuhan |
| 29544 | 35 | D | 3 | Spain |
| 29593 | 2 | I | 2 | USA |
| 29686 | 1 | I | 7 | Iceland, England, Thailand |
| 29726 | 1 | D | 2 | England |
| 29730 | 37 | D | 2 | England |
| 29730 | 30 | D | 2 | England, Poland |
| 29756 | 7 | D | 6 | India, England, USA, Netherlands, Canada |
| 29759 | 5 | D | 3 | USA |
| 29760 | 2 | D | 4 | USA |
| 29788 | 2 | D | 3 | England |

## ACKNOWLEDGEMENTS

Figures were created using Biorender.com. The full acknowledgements for the GISAID sequence providers can be found in Table S2. We thank the NSF GRFP, Stanford Bio-X, and Stanford Precision Health and Integrated Diagnostics Center (PHIND) for funding.

